# Dehydration bouts prompt increased activity and blood feeding by mosquitoes

**DOI:** 10.1101/120741

**Authors:** Richard W. Hagan, Elise M. Szuter, Andrew E. Rosselot, Christopher J. Holmes, Samantha C. Siler, Andrew J. Rosendale, Jacob M. Hendershot, Kiaira S. B. Elliott, Emily C. Jennings, Alexandre E. Rizlallah, Yanyu Xiao, Miki Watanabe, Lindsey E. Romick-Rosendale, Jason L. Rasgon, Joshua B. Benoit

**Affiliations:** Department of Biological Sciences, University of Cincinnati, Cincinnati, OH 45221 USA.; Department of Mathematical Sciences, University of Cincinnati, Cincinnati, OH 45221 USA.; Division of Pathology, Cincinnati Children’s Hospital Medical Center, Cincinnati, OH 45229 USA.; Department of Entomology, Pennsylvania State University, Center for Infectious Disease Dynamics and Huck Institutes for Life Sciences, Pennsylvania State University, University Park, PA 16802 USA.

**Keywords:** blood feeding, dehydration, mosquitoes, landing activity

## Abstract

Mosquitoes are prone to dehydration and respond to this stress through multiple mechanisms, but previous studies have examined very specific responses and fail to provide an encompassing view of the role that dehydration has on mosquito biology. This study examined underlying changes in biology of the northern house mosquito, *Culex pipiens*, associated with short bouts of dehydration. We show that dehydration increased blood feeding propensity of mosquitoes, which was the result of both enhanced activity and a higher tendency to land on a host. Mosquitoes exposed to dehydrating conditions with access to water or rehydrated individuals experience no water loss and failed to display behavioral changes. RNA-seq and metabolome analyses following dehydration indicated that factors associated with energy metabolism are altered, specifically the breakdown of trehalose to yield glucose, which likely underlies changes in mosquito activity. Suppression of trehalose breakdown by RNA interference reduced phenotypes associated with dehydration. Comparable results were noted for two other mosquito species, suggesting this is a general response among mosquitoes. Lastly, field-based mesocosm studies using *C*. *pipiens* revealed that dehydrated mosquitoes were more likely to host feed, and disease modeling indicates dehydration bouts may increase transmission of West Nile virus. These results suggest that periods of dehydration prompt mosquitoes to utilize blood feeding as a mechanism to obtain water. This dehydration-induced increase in blood feeding is likely to intensify disease transmission during periods of low water availability.

**Significance:** Dehydration stress has substantial impacts on the biology of terrestrial invertebrates. To date, no studies have elucidated the difference between dehydration exposure and realized water loss in relation to mosquito behavior and physiology. Our experiments show that direct dehydration stress increases mosquito activity and subsequent blood feeding, likely as a mechanism to locate and utilize a bloodmeal for rehydration. These dehydration-induced phenotypes were linked to altered carbohydrate metabolism that acts as a source of energy. This study provides important insight into the impact of mosquito-dehydration dynamics on disease transmission that is likely general among mosquitoes.

## Introduction

Dehydration significantly impacts the geographical distribution, reproductive capacity, and longevity of terrestrial arthropods^1-4^. With more, and extended, periods of drought predicted to occur as climate change progresses^5,6^, the ability to tolerate bouts of dehydration is becoming increasingly critical to insect survival. Water loss for terrestrial invertebrates occurs through excretion, respiratory gas exchange, and cuticle transpiration^7,8^. Insects residing in dry environments significantly minimize water loss by mechanisms such as suppressing rates of cuticular water loss and reducing excretion^8^. If water balance cannot be maintained, a complement of mechanisms is employed to prevent cellular damage from osmotic stress. These changes include increased expression of heat shock proteins^3,8-10^, shifts in osmoprotectant metabolites^9-12^, and differential regulation of aquaporins to alter the movement of water between the hemolymph and intracellular components^13-15^.

Dehydration tolerance in mosquitoes is a significant factor in determining their ability to expand to new regions and also dictates their survival under periods of low water availability^7,16-20^. For mosquitoes that transmit human malaria, there are many recent studies examining molecular and physiological changes associated with dehydration or dry-season conditions^19-21^. Rearing *Anopheles gambiae* under dry season conditions shifted their metabolite levels, suppressed reproduction, and altered spiracle size^18-23^. Microarray analyses of dehydrated *A. gambiae* revealed increased expression of genes associated with stress response and DNA repair^21^. Although these studies have identified many interesting aspects associated with the dynamics between dehydration and mosquito physiology, none have examined the effects of dehydration bouts through the utilization of integrated studies across multiple aspects of mosquito biology.

In this study, we examined the effect of short bouts of dehydration on the physiology of mosquitoes via RNA-seq analyses and by blood feeding and physiological assays. Our initial hypothesis was that dehydration would yield a suppression of metabolism, reduced activity, and lower levels of blood feeding for the northern house mosquito, *Culex pipiens*. However, we found that short periods of dehydration prompted an increase in transcript levels of genes associated with energy metabolism, increased activity, elicited a higher propensity for mosquito host landing, and elevated blood feeding. Offering a source of drinking water during dehydration or rehydration inhibited these phenotypic changes, suggesting that dehydration, rather than exposure to low relative humidity (RH), prompts the observed phenotypes. Similar results were noted for two other mosquito species, *Aedes aegypti*, and *Anopheles quadrimaculatus*, suggesting that these findings are generalizable across mosquito species. Mesocosm-based studies indicated these dehydration-induced responses also occur under simulated field conditions. Furthermore, modeling of our findings provide support for the hypothesis that increased blood feeding, due to dehydration, will likely alter disease transmission. Our study indicates that brief bouts of dehydration substantially alters the physiology and subsequent behavior of mosquitoes in a manner that has significant implications for mosquito-environment-blood feeding dynamics.

## Results

### Shifts in activity and blood feeding after dehydration stress

During initial dehydration experiments, we observed that dehydrated mosquitoes appeared more active compared to control, non-dehydrated counterparts. To test this observation, total activity was assessed for *C. pipiens* females under dehydrating (75% RH) and non-dehydrating (100% RH) conditions. Exposure to dehydration increased activity after 30 hours (∼15-20% water loss, Fig. S2), and activity levels were elevated until dehydration-induced mortality occurred at a loss of ∼35% water content (Fig. 1A). In conjunction with increased activity, the propensity of mosquitoes to land on a host mimic (a heated, membrane covered blood feeder, Hemotek, UK) increased nearly 4- to 5-fold after losing 15-20% of water content at 75% RH, but not when held under non-dehydrating conditions (100% RH) or at 0% RH with access to liquid water for ingestion (Fig. 1B). This indicates that dehydration stress, not dry conditions, was responsible for the observed phenotypes. Comparative studies using *A. aegypti* and *A. quadrimaculatus*, confirmed that dehydration induced increases in both activity and host landing is likely a general response across diverse mosquito species (Fig. S1-S3). Lastly, the propensity of *C. pipiens* to ingest a blood meal increased significantly after a 10-15% loss in water and was the highest when water loss was 20-30% (Fig. 1C). When temperature and relative humidity were examined in relation to dehydration levels, the dehydration threshold of 20% can be reached within as little as six-eight hours at low (Fig. 1D), but ecologically relevant, RHs (60-80%) and typical summer temperatures (26-28°C). These results indicate that dehydration bouts increase the propensity of females to ingest a blood meal and this impact can be exacerbated by ecologically-relevant temperature and RH dynamics.

**Fig.1.**
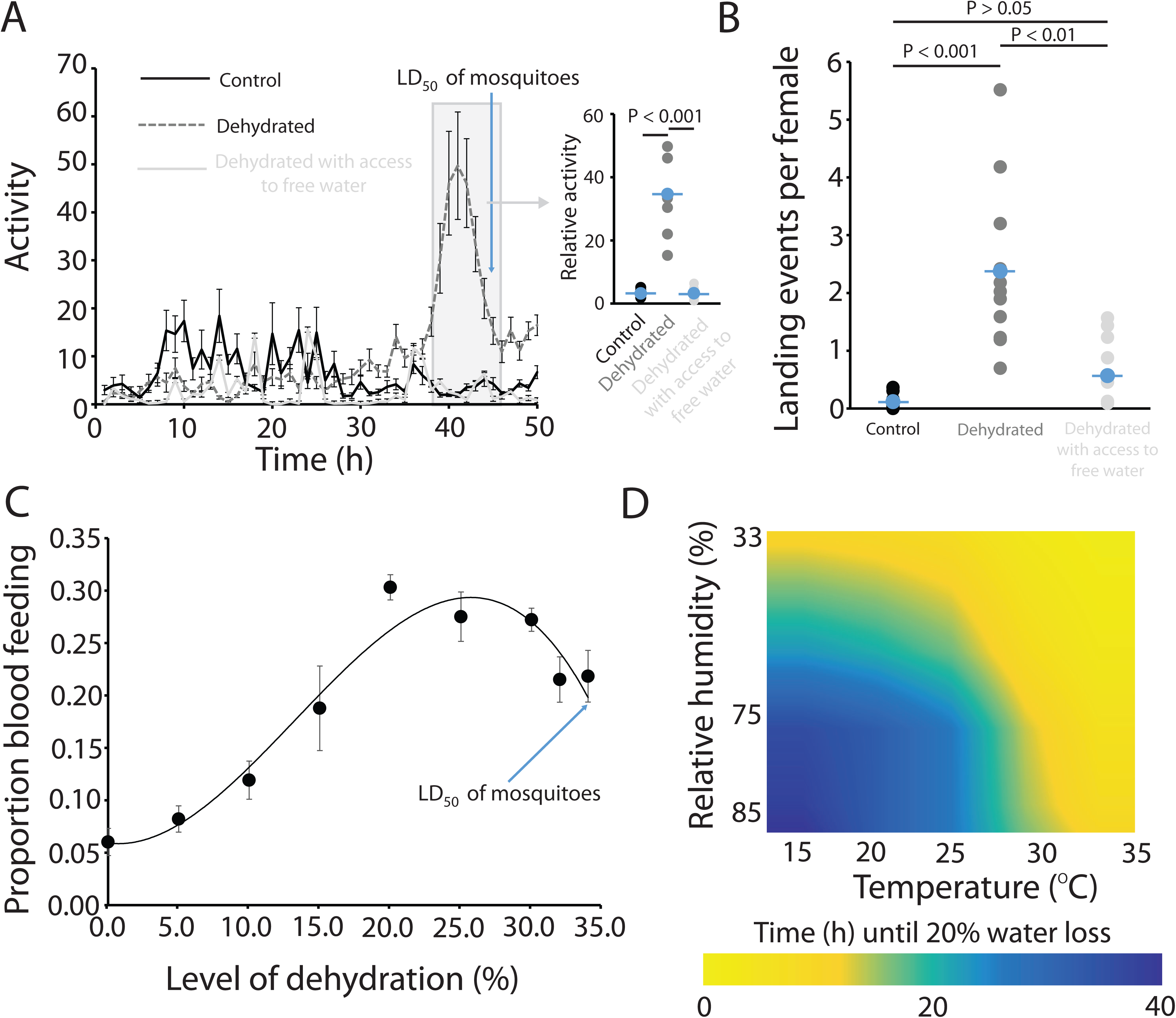
Dehydration increases mosquito activity and blood feeding. (*A*) Activity measured by a Locomotor Activity Monitor 25 (LAM25) system during the course of dehydration. Inset represents combined results from 38-44 hours. Mean ± SE represents 48 mosquitoes. (*B*) Number of landing events per mosquito over the course of one hour. Mean ± SE represents 12 independent replicates of 30-40 mosquitoes. (*C*) Proportion of mosquitoes that blood fed within two hours of host availability. Mean ± SE represents 10 replicates of 50 mosquitoes. (*D*) Time until the 20% dehydration point at varying temperatures and relative humidity (RH). Three replicates of 40 mosquitoes were conducted at each RH (33, 75, and 100%) at each temperature. Statistical analyses were conducted by a t-test or one- and two-way ANOVA followed by Tukey’s HSD post-hoc analysis where appropriate.

### Transcriptional and metabolomic changes associated with *C. pipiens* dehydration

To investigate the underlying mechanisms associated with the dehydration-induced phenotype, we utilized a combined RNA-seq and metabolomic approach. Dehydration exposure (20% loss of water) yielded over 1200 genes with differential expression compared to fully hydrated individuals, a subset of which was validated through quantitative PCR (Fig. 2A, Fig. S4, Table S1). Enrichment was noted for Gene Ontology terms associated with carbohydrate metabolism (Fig. 2B, Table S2). Metabolomic analyses revealed a substantial increase in the levels of glucose during dehydration stress, supporting our RNA-seq data (Fig. 2C). Trehalose content declined by over 60% and is likely the source of the increase in glucose (two glucose molecules are generated per breakdown of one trehalose). Based on these results, we hypothesize that altered carbohydrate metabolism, specifically the breakdown of trehalose to glucose, may underlie the phenotypic changes following dehydration in *C*. *pipiens*.

**Fig.2.**
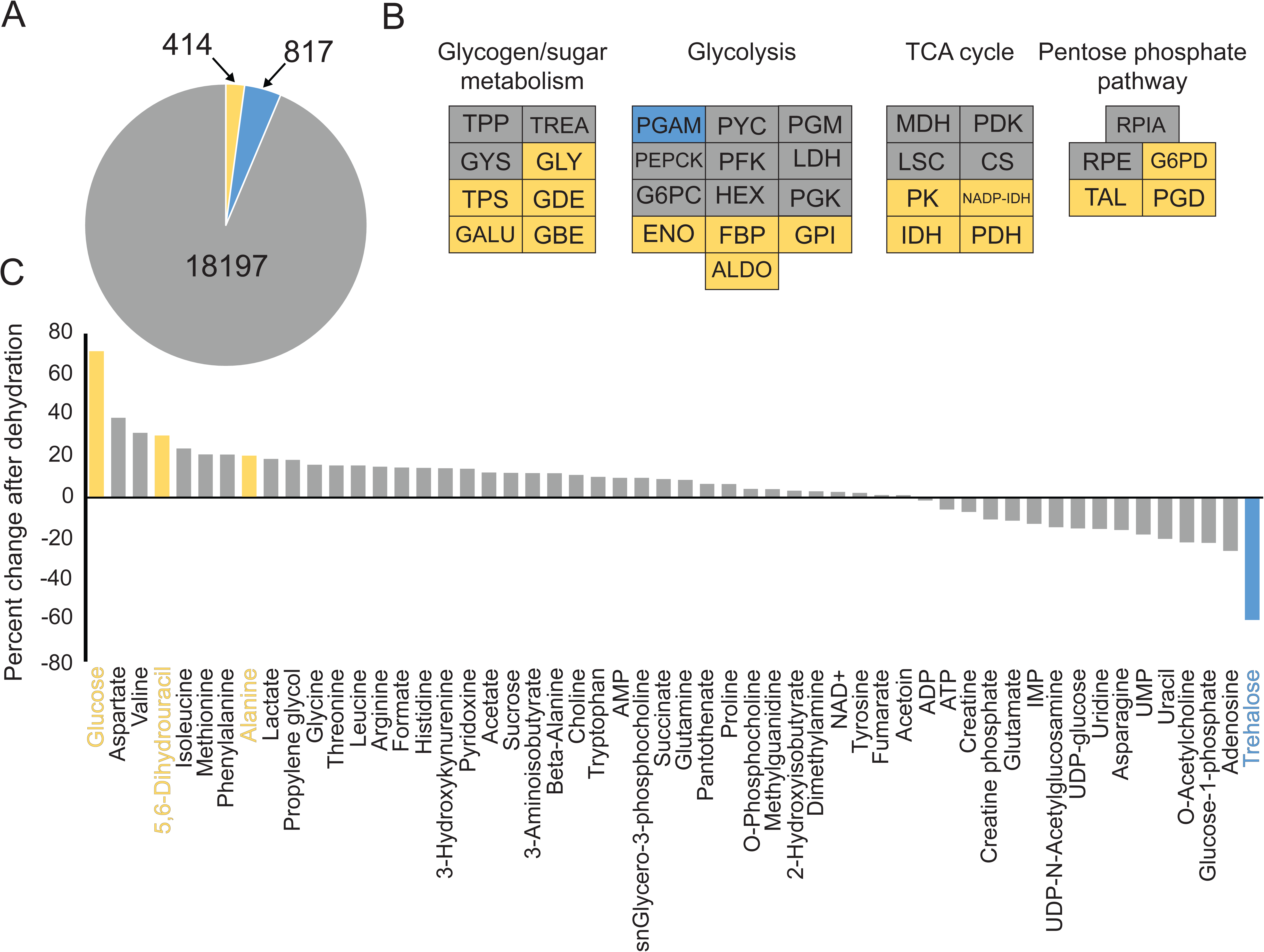
Dehydration enriched carbohydrate metabolism prompts a trehalose to glucose shift. (*A*) RNA-seq analyses following dehydration yielded differential expression of 1231 genes. (*B*) Increased expression for multiple carbohydrate-associated genes. (*C*) Metabolomics following dehydration stress. In (A), (B) and (C), blue denotes increased levels during dehydration, gray indicates no difference, and yellow is decreased during dehydration compared to hydrated individuals.

### Trehalose to glucose conversion during dehydration and suppression of trehalose metabolism

Spetrophotometric assays validated our metabolomics results by showing glucose increased and trehalose decreased following dehydration exposure when measured (Fig. 3A). The increase of glucose and decrease of trehalose was augmented under slow dehydration, suggesting that the rate of dehydration has a substantial impact on dehydration-induced phenotypes (Fig. 3A). Glycogen levels in response to dehydration were examined since this reserve serves as a major source of trehalose (Fig. S5). We noted differences between control and dehydrated mosquitoes under slow dehydration (75% RH), which have a significant, albeit slight, decline in glycogen (Fig. S5). These findings correlate with those from previous studies, which showed that slow compared to rapid dehydration induces more pronounced physiological and behavioral responses^10^. Mosquitoes held under dehydrating conditions, but with access to free water, showed no reduction in glycogen levels compared to those that experienced dehydration stress without water (Fig. S5). Starvation does not serve as a major contributor to our observed phenotypes since mosquitoes held at higher relative humidity, or allowed access to free water, survive for nearly 4-5 days before starvation-induced death compared to dehydrated individuals (*p*<0.0001, log-rank test; Fig. S6). These results suggest that a bout of dehydration alters trehalose and glycogen metabolism, thus generating more glucose. This phenotype is augmented when dehydration occurs at a slower rate.

**Fig.3.**
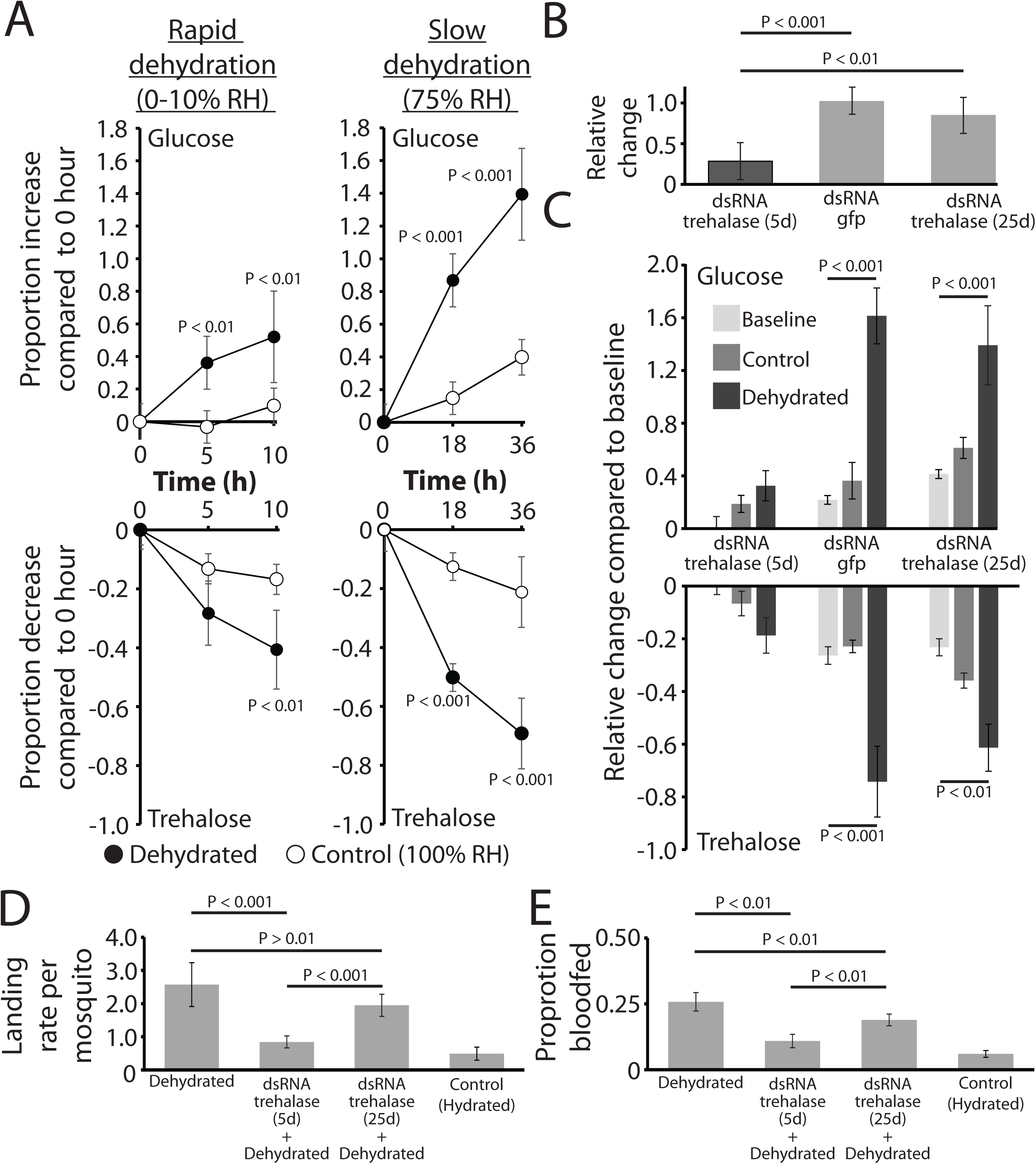
Glucose and trehalose shifts directly impact mosquito blood feeding-dehydration dynamics. (*A*) Rapid and slow dehydration differentially altered levels of glucose and trehalose. Mean ± SE for 10 replicates of samples from four-five mosquitoes at each time point. (*B-C*) Knockdown of *trehalase* prevented the trehalose to glucose conversion during dehydration. Mean ± SE for 12 mosquitoes at each time point. (*D-E*). Landing on the host (mean ± SE for 8 replicates at each time point) and blood feeding (mean ± SE for 8 replicates at each time point) changed following *trehalase* suppression and dehydration. Statistical analyses were conducted by a one- or two-way ANOVA followed by Tukey’s HSD post-hoc analysis where appropriate.

We observed that trehalose catabolism was augmented during slow dehydration, leading us to hypothesize that the rate of dehydration has a substantial impact on dehydration-induced phenotypes through the breakdown of trehalose. To link metabolism of trehalose with shifts in blood feeding, we utilized RNA interference to suppress the levels of *trehalase*, the enzyme involved in the conversion of trehalose to glucose, during the course of dehydration. dsRNA injection targeting *trehalase* resulted in nearly an 80% reduction in transcript levels compared to control mosquitoes (Fig. 3B). *Trehalase* transcript levels recovered 25 days after dsRNA injection (Fig. 3B). When trehalose and glucose were measured following *trehalase* knockdown, there was a slight, non-significant, trehalose decrease and glucose increase following dehydration stress (Fig. 3C). This lack of trehalose to glucose conversion after *trehalase* suppression was in stark contrast to shifts in trehalose and glucose observed when mosquitoes were injected with dsRNA targeting *green fluorescent protein* (*gfp*) or when the impact of dsRNA targeting *trehalase* had dissipated after 25 days (Fig. 3C). The proportion of mosquitoes that landed on a host mimic following dehydration was reduced by over 50% after knockdown of *trehalase* and the response was partially recovered 25 days after dsRNA injection (Fig. 3D). Similarly, increased blood feeding following dehydration was suppressed following knockdown of *trehalase* and recovered once the dsRNA effects began to decline after 25 days (Fig. 3E). Thus, the inhibition of trehalose to glucose conversion during dehydration inhibits the dehydration-induced phenotype of increased host interactions.

### Mesocosm-based analyses and modeling of disease transmission

To assess if dehydration status impacted blood feeding under field conditions, we designed a mesocosm-based study to examine water content of mosquitoes attempting to feed on a Hemotek-based host mimic. Briefly, *Culex pipiens* adults were allowed to emerge within an outdoor mesh-enclosed mesocosm conducted in June 2016 at the University of Cincinnati Center for Field Studies and were provided free access to water and sugar. After eight-twelve days, mosquitoes were allowed access to an artificial host and individuals that landed and probed during the first 60 minutes were collected. In addition, we collected ∼10-20 mosquitoes at the completion of the experiments that did not respond to the host mimic. We performed these studies under two specific weather conditions related to water availability; precipitation had (wet) or had not (dry) occurred in the last 24 hours. Mosquitoes that more frequently landed on a host mimic had reduced water content in comparison to those that did not respond to the host mimic (Fig. 4A). The effect of hydration status on host landing was exacerbated in dry conditions during the preceding 24 hours compared with wet conditions (Fig. 4A). These results indicate that our laboratory-based studies related to mosquito dehydration exposure are relevant under ecological conditions.

**Fig.4.**
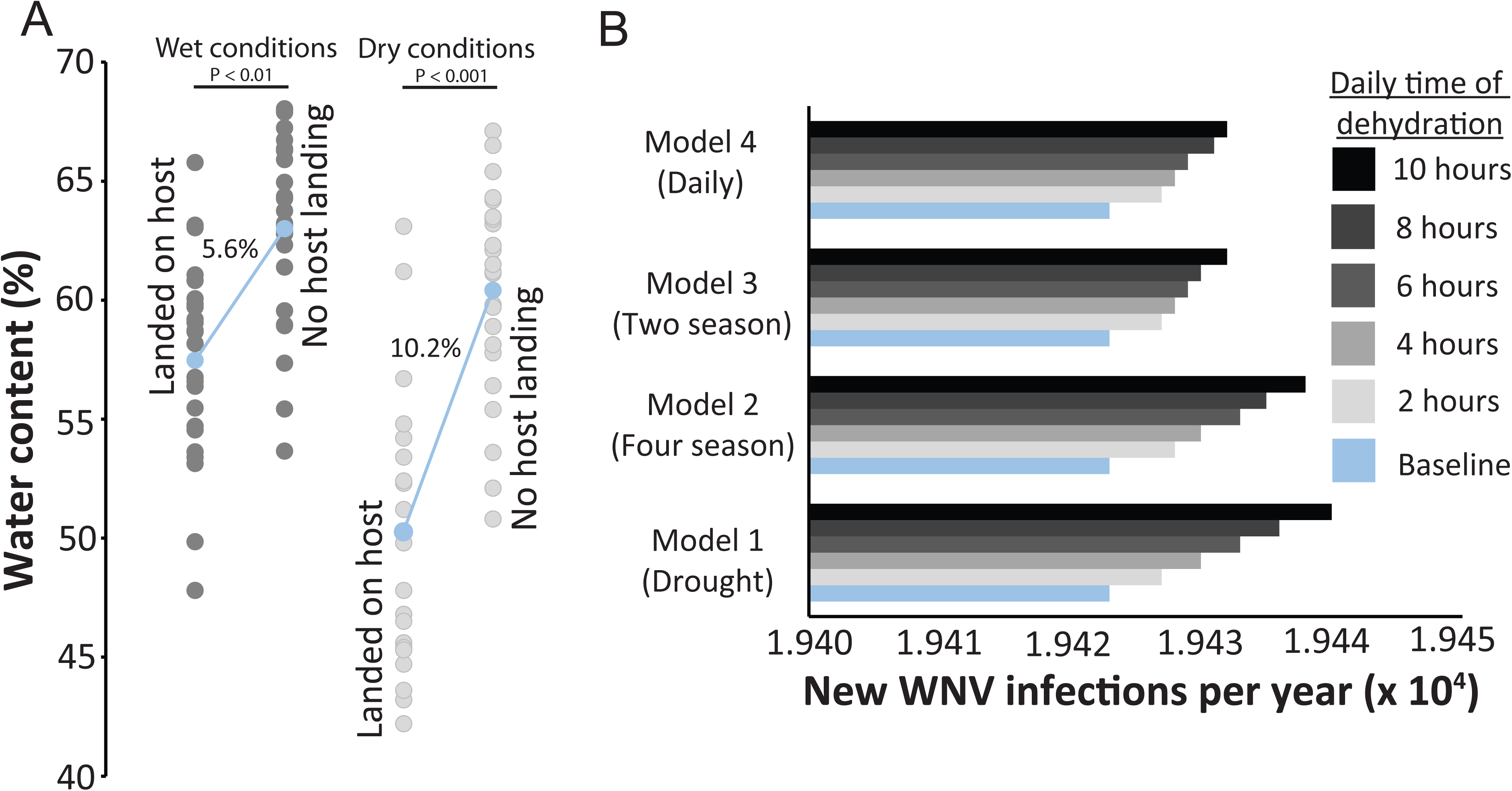
Dry field conditions increase host landing and impact disease transmission based on West Nile virus transmission modeling. (*A*) Water content assessment of feeding mosquitoes following 24 hours with and without precipitation. Mean ± SE for 8 replicates at each time point. (*B*) Predicted transmission of WNV will increase as the result of increased blood feeding due to dehydration exposure. Modeling described in Supplemental Methods 1.

Lastly, to determine if the increased propensity to blood feed results in the potential to prompt higher disease transmission, we incorporated the impact of short dehydration bouts into a disease transmission model (Fig. 4; Supplemental Materials 1). Herein we demonstrate that *C. pipiens* blood feed more frequently following dehydration (Fig. 1), which we subsequently confirmed using different dehydration and rehydration protocols (Fig. S7). This allowed comparable blood feeding rate to be measured for dehydration, control, and rehydrated mosquitoes. Blood feeding rates were converted to a biting rate based on proportional changes in blood feeding according to previous studies^24-26^. With these varying biting rates, we developed four models that incorporate the increased biting rate due to daily dehydration, multiple bouts of dehydration, dehydration in relation to seasonal changes, and the impact of multiple seasons into the potential transmission of West Nile virus (Fig. 4B). All models predicted that dehydration bouts yielded gradual increases in disease transmission with the largest increase in potential disease transmission after a prolonged period (10 hours) of dehydration.

## Discussion

Mosquitoes must contend with a variety of environmental conditions, and dehydration is one of the most critical factors influencing their distribution and survival. This study revealed that bouts of dehydration alter the physiology and behavior of mosquitoes, yielding increased activity and rate of blood feeding, when compared to fully hydrated individuals. This shift in activity and blood feeding will be manifested in an increase in the transmission of mosquito-borne diseases following bouts of dehydration. We have provided molecular evidence that the breakdown of trehalose into glucose plays a central role in promoting the dehydration-induced phenotype in mosquitoes. Taken together, these findings indicate that mosquito hydration can drastically shift mosquito phenotypes. Thus, this information must be accounted for in studies that assess potential disease transmission.

Many recent studies have assessed the impact of dehydration on mosquito physiology^18-20^ and more specifically the distinct changes in many amino acids and sugars due to dehydration exposure^18-20^. The increase in glucose levels following dehydration observed in *An. gambiae* females after dehydration^19^ was similar to the increase seen in this study. Furthermore, water deficit-based fluctuations in sugars and polyols have been documented in many insect systems^11,27,28^. These changes likely prevent additional body water loss by altering the colligative properties of hemolymph and may reduce potentially damaging interactions among biological molecules. Along with the conversion of trehalose to glucose, we noted a significant, albeit small, decline in glycogen following prolonged dehydration, which yields four-five units of water for each unit of glycogen metabolized^29^. This information indicates that increased glucose levels are a common physiological response following dehydration for many insect systems.

Other than the putative protective and water generation roles for the increased glucose during dehydration, there have not been other roles established for the glucose shift. Here we show that suppression of metabolic pathways necessary to generate glucose prevented dehydration-induced behaviors. Increased activity, host landing, and blood feeding following dehydration are subsequently recovered once the temporal effects of dsRNA targeting *trehalase* have faded. Previous studies have demonstrated that increased carbohydrate oxidation, along with elevated levels of constituents of the TCA cycle, are critical in providing fuel for wing muscles during flight^30-32^. In A. *gambiae*, expression of genes associated with glycogenolytic processes increase when mosquitoes are exposed to dry season conditions^19^. This phenomenon is comparable to our observed results in *C. pipiens*. This suggests that increased levels of sugar and amino acids may act beyond their role in stress protection and act as an energy source to support behavioral shifts that will increase blood feeding to enable hydration.

The rate of dehydration is influenced by both the temperature and relative humidity (RH), where high temperature and low RH yield the highest water loss rates. Under conditions that induce dehydration rapidly (33% RH, 35°C), water loss reached a critical threshold of 20% within 9 hours. Less severe conditions (75% RH, 25°C) reduced this threshold to approximately ∼24-28 hours. Importantly, the most drastic effects on glucose concentration, activity, and blood feeding occurred during slower dehydration, suggesting that the findings from this study are ecologically relevant, and will be critical in the analysis of the dehydration-induced phenotypes. These findings were supported under field conditions, where individuals with lower water content, indicative of dehydration, were more likely to land on a host. Lastly, WNV transmission modeling revealed that increased blood feeding will increase number of new infections, suggesting our results need to be accounted for in future studies related to modeling viral transmission by mosquitoes.

In conclusion, bouts of dehydration alter carbohydrate metabolism, which in turn boosts mosquito activity and prompts increased blood feeding. This blood feeding is not the direct result of starvation, but rather likely serves as a mechanism for immediate rehydration. During dry periods it is likely that rehydration through host feeding could be more feasible than ingestion of freestanding water, since temporal water pools may be difficult to locate in some environments relative to a host. This increased propensity of blood feeding will likely increase the potential for disease transmission, due to a higher biting rate during dry periods with low precipitation. Recent studies have indicated that the transmission rates of West Nile virus are impacted by dry periods, where drought has been identified as one of the most critical factors underlying WNV outbreaks^33-38^. Specifically, there have been four proposed mechanisms that account for why dehydrating conditions may impact disease transmission dynamics. First, periods of drought likely bring mosquitoes and animal hosts in close contact due to the scarcity of water sources^39^. Second, dry conditions will concentrate nutrients in water pools^34,35^, providing the resources necessary for larval development. Third, periodic drying will suppress levels of predators and other competitors and allow mosquitoes to proliferate^40^. Fourth, lack of rain will allow for water to pool in storm drains, which yield temporal water pools for mosquito development^33-35^. In this study, we establish a fifth major dehydration-associate mechanism that will impact disease transmission – dehydration-induced shifts in physiology and behavior of mosquitoes prompting increased blood feeding. This shift in biting rate, due to dehydration, must be accounted for in studies that seek to predict the prevalence and potential outbreaks of mosquito-borne diseases such as West Nile, Zika, Chikungunya, and Dengue viruses.

## Materials and Methods

### Mosquito husbandry

Mosquitoes were maintained in a climate-controlled facility at the University of Cincinnati. Ambient temperature was held between 21-28°C and relative humidity (RH) was held between 70-80%. A 12-12 h light-dark photoperiod was maintained for the duration of the experiment. For each species, eggs were harvested after oviposition. Postemergence, larvae were separated into tanks of 75-85 individuals per 0.5 L water to control for density during maturation. Larvae were fed finely ground fish food (Tertramin) with added yeast extract (Difco). Adults were kept in 12” × 12” × 12” mesh cages with unrestricted access to clean water and a 10% sucrose solution. Mosquitoes used in this study were 10-14 days post adult emergence. Three species were used in this study: *Aedes aegypti* Benzon, *Culex pipiens* Buckeye^41^, and *Anopheles quadrimaculatus* Benzon.

### Dehydration experiments

Mosquitoes were dehydrated by placing groups of 20 mosquitoes in a 50 ml centrifuge tube covered with mesh. These tubes were placed at 0% or 75% RH and predominantly removed after either 10-15% water loss (18-20 hours at 75% RH or 4-6 hours at 0% RH) or 20-25% water loss (36-40 hours at 75% RH or 9-11 hours at 0% RH). Control mosquitoes were held at 100% RH. Individuals exposed to dehydrating conditions (75% RH, generated with saturated NaCl solutions) and allowed access to free water remained fully hydrated. In addition, a subset of dehydrated mosquitoes were moved to 100% RH (generated with deionized water) with access to free water to determine if the phenotypes induced by dehydration can be reversed. A schematic of the dehydration experiment are included as Figure S8. This allowed comparisons among dehydrated, fully hydrated, and fully hydrated under dehydrating conditions to elucidate the effect of body water content reduction versus exposure to dehydrating conditions, which has been lacking in other studies on the response of arthropods to dehydration.

### RNA-seq, metabolomic analyses, and gene suppression

RNA-seq analyses were conducted on dehydrated (20% loss) and fully hydrated mosquitoes. RNA was extracted with Trizol based upon manufacturers’ methods (Invitrogen). Two independent biological libraries were prepared and sequencing was conducted at the Cincinnati Children’s Hospital Medical Center RNA Sequencing Core. Reads were mapped to predicted genes from the *Culex quinquefasciatus* genome (Gene set version 2.1) using CLC Genomics (CLC Bio) based upon methods used in Benoit *et al*.^42^ and Rosendale *et al*.^28^ Differentially expressed genes were determined using TPM with P values corrected using the Bonferroni method to reduce false discovery. Gene Ontology (GO) terms were identified with Blast2GO^43^ and g:Profiler^44^. Specific KEGG pathways were examined for enrichment through the use of GSEA^45,46^. *Trehalase* levels were suppressed through the use of dsRNA injection (1.0-1.5 μg/μl), which was generated and injected according to methods developed previously^9^. qPCR was utilized to validate RNA-seq and *trehalase* knockdown and conducted according to previous studies^28^ (Table S3).

Metabolomics were conducted on mosquitoes after 18-20 hours of dehydration at 75% RH (Dehydrated) or 100% RH (Control) according to methods used by Rosendale *et al*.^28^. Briefly, mosquitoes were collected and dried to establish the dry mass, which was used to standardize each sample. Eight replicates of thirty to forty mosquitoes were collected and analyzed for the control and dehydrated samples.

### Survival analyses

Mosquito survival assays were conducted to determine the time until death. Three conditions were examined, dehydration, dehydration-inducing conditions with access to free water, and exposure to high RH (100% RH) under similar conditions as listed above. Mosquito death was determined when mosquitoes failed to respond to stimulus and move. Differences in survival were assessed based on the LD_50_ determined with a Kaplan-Meier survival analyses (JMP). Three groups of 50 mosquitoes were used for each treatment.

### Nutrient reserves

Levels of mosquito nutrient reserves were measured using spectophotometric methods according to previous studies^1,47-50^. Briefly, three mosquitoes were combined and homogenized with the use of a bead blaster in 250 μl of 2.0% Na_2_SO_4_. Soluble sugars were extracted using a 1:1 chloroform/methanol mixture. Glycogen was quantified with an anthrone assay^49^. Trehalose and glucose assays were conducted as described in Liu *et al* ^51^.

### Landing and blood-feeding assays

The number of mosquitoes that landed on the host was determined based on techniques modified from Barnard *et al*.^52^. Twenty to thirty mosquitoes were released into a 60 × 60 × 60 cm cage (Bugdorm) and allowed two hours to equilibrate. An artificial host (Hemotek) was covered three times with Parafilm, filled with water, heated to 37°C, and attached to the top of the cage. Mosquitoes that landed and remained on the artificial host for at least 5 seconds were used to differentiate incidental contact from exploratory (foraging) contact. The number of mosquitoes was counted at five-minute intervals for sixty minutes. Blood feeding was measured using the same apparatus and setup except the Parafilm was pulled thin to allow successful probing and the feeding disk was filled with chicken blood (Pel-Freeze).

### Activity assays

General activity of mosquitoes was assessed using a Locomotor Activity Monitor 25 (LAM25) system (TriKinetics, Waltham, MA) based on methods previously developed for mosquitoes^53^. Briefly, females 7-10 days post-emergence (mated, but not blood fed) were moved into 25 × 150 mm clear glass tubes. Flight activity was recorded as the number of times a mosquito passed across an infrared beam. All recordings were conducted under continual light cycles to prevent confounding effects of light:dark transition on the dehydration response, which has been noted in other studies ^53,54^. Three groups were analyzed: mosquitoes held at 75% RH (dehydrated), those held at 100% RH (fully hydrated), and mosquitoes at 75% RH with access to free water (fully hydrated, but exposed to dehydration-inducing conditions).

### Field-based mesocosm studies and modeling

Lab reared mosquitoes were released into a 6’ × 6’ × 6’ mesh-covered cage (Bioquip) at the University of Cincinnati Center for Field Studies (39.285N, 84.741W). Water and 10% sucrose solution were provided *ad libitum* and changed every two days. After five-six days, a host mimic was introduced at dusk and mosquitoes that landed on the host mimic (Hemotek) were collected and immediately frozen. In addition, eight-ten mosquitoes that did not land were collected with an aspirator. Water content of these mosquitoes was determined according to Benoit and Denlinger^16^.

Modeling of WNV transmission was developed based on previous studies^24-26^. We modified the biting rates in each model based upon proportional changes in blood-feeding that examined the impact of dehydration, dehydration with access to water, and those experiencing no dehydration stress on blood feeding (Fig. S7). Specific details of the modeling are included in the supplemental methods (Supplemental Materials 1).

### Statistics and replication

Replicates throughout the experiments are biologically independent and distinct samples. Sample sizes are listed in the methods section or the appropriate figure legend. Statistical significance between treatments and controls is indicated within each figure and/or in the figure legend. Statistical tests are listed within the respective figure legends. All statistical analyses were performed using JMP version 11 (SAS) or R-based packages.

## ACKNOWLEDGEMENTS

We appreciate the helpful comments on this manuscript from David L. Denlinger (Ohio State University) and Brian L. Weiss (Yale University). This work was supported in part by a Faculty Development Grant from the University of Cincinnati (J. B.), Ohio Supercomputer Center (J. B.), United State Department of Agriculture’s National Institute of Food and Agricultural Grant (2016-67012-24652, A.R.), NMR-Based Metabolomics Core at the Cincinnati Children’s Hospital Medical Center, and National Institutes of Health (R01AI116636 and R21AI128918, J.R.).

## Table S1.

List of genes with differential expression following exposure to dehydration compared to fully hydrated individuals.

## Table S2.

Differential expression of genes associated with carbohydrate metabolism.

## Table S3.

Quantitative PCR and dsRNA primers.

## Supplemental Figures

### Fig. S1.

Activity through the course of dehydration for *Culex pipiens, Aedes aegypti*, and *Anopheles quadrimaculatus*. Each time course represents the mean ± SE represents 48 mosquitoes. Statistical analyses were conducted by a t-test. Specific shaded areas are highlighted by individual plots to allow comparison between different periods of dehydration.

Dehydration status of mosquitoes after 40 hours at 75% RH. Statistical analyses were conducted by a one-way ANOVA or t-test followed by Tukey’s post-hoc analysis.

### Fig. S3.

Number of landing events per mosquito over the course of one hour. Mean ± SE represents 11-13 independent replicates of 30-40 mosquitoes. Statistical analyses were conducted by a one- or two-ANOVA followed by Tukey’s HSD post-hoc analysis.

### Fig. S4.

RNA-seq validation by qPCR. Each qPCR measurement was replicated four times. RNA-seq and qPCR results were compared with a Pearson’s Correlation Coefficient.

### Fig. S5.

Glycogen content for *Culex pipiens* when held at 75% relative humidity (RH), 100% RH, and 75% RH with free access to water. Mean ± SE for 8 mosquitoes at each time point. Statistical analyses were conducted by a one- or two-way ANOVA followed by Tukey’s HSD post-hoc analysis

### Fig. S6.

Survival of mosquitoes when held at 75% relative humidity (RH), 100% RH, and 75% RH with free access to water. Mean ± SE for 3 replicates of 50 mosquitoes at each time point. Differences in survival were assessed through a probit analysis.

### Fig. S7.

Proportion of mosquitoes (*Culex pipiens*) blood feeding held under different dehydration, rehydration, and dehydrating conditions with access to water protocols. Statistical analyses were conducted by ANOVA followed by Dunnett’s post-hoc analysis. *, *p*< 0.05.

### Fig. S8.

Schematic for dehydration exposures illustrating both dehydration and rehydration protocols. Each dehydration/rehydration point was not used for all assays.

